# Topographical cuescontrol the morphology and dynamics of migrating cortical interneurons

**DOI:** 10.1101/452359

**Authors:** Claire Leclech, Marianne Renner, catherine Villard, Christine Métin

## Abstract

In mammalian embryos, cortical interneurons travel long distances among complex three-dimensional tissues before integrating into cortical circuits. Several molecular guiding cues involved in this migration process have been identified, but the influence of physical parameters remains poorly understood. In the present study, we have investigated *in vitro* the influence of the topography of the microenvironment on the migration of primary cortical interneurons released from mouse embryonic explants.

We found that arrays of 10 μm-sized PDMS micro-pillars, either round or square, influenced both the morphology and the migratory behavior of interneurons. Strikingly, most interneurons exhibited a single and long leading process oriented along the diagonals of the square pillared array, whereas leading processes of interneurons migrating in-between round pillars were shorter, often branched and oriented in all available directions. Accordingly, dynamic studies revealed that growth cone divisions were twice more frequent in round than in square pillars. Both soma and leading process tips presented forward directed movements within square pillars, contrasting with the erratic trajectories and more dynamic movements observed among round pillars. In support of these observations, long interneurons migrating in square pillars displayed tight bundles of stable microtubules aligned in the direction of migration.

Overall, our results show that micron-sized topography provides global spatial constraints promoting the establishment of two different morphological and migratory states. Remarkably, both states belong to the natural range of migratory behaviors of cortical interneurons, highlighting the potential importance of topographical cues in the guidance of these embryonic neurons, and more generally in brain development.

## INTRODUCTION

Neuronal migration is a major phase of nervous system development. Migration alterations are often associated with brain disorders like microcephaly or lissencephaly [1-3]. Among the embryonic neurons, cortical interneurons travel over long distances in the developing brain, from the ventral ganglionic eminences where they are generated to the dorsally located cortex, where they play a crucial role in modulating the activity of cortical circuits in the adult [4-6]. Within the embryonic cortex, migrating interneurons first follow tangential routes that disperse them in the whole cortical domain and then reorient radially to colonize the cortical plate [7-10], see reviews in [11-13]. During migration, interneurons exhibit changing morphologies due to the stochastic branching of their leading process [14]. Their reorientation away from the tangential pathways mostly relies on the stabilization of novel branches with an appropriate orientation [7,9,10,15,16].

The factors influencing interneuron migration have been extensively studied both *in vivo* and *in vitro,* mainly focusing on the cellular and intracellular responses to molecular guidance cues. In the current view, the expression of repulsive factors in the ventral regions combined with the presence of attractive and motogenic signals in the dorsal cortex play a central role in the guidance of interneurons [17-28] reviewed in [5,11]. Migrating interneurons also establish adhesive interactions with the extracellular matrix and cellular components distributed on their pathways [29-31]. They are moreover exposed to various topographic and mechanical cues linked to the properties of their extracellular environment including various cellular substrates, e.g. neuronal progenitors, corticofugal axons or post-mitotic neurons [32-35].

There is growing evidence indicating that physical parameters can have a crucial influence in the development of the nervous system [36,37]. It has been shown for example that the developing brain displays regional and temporal variations in stiffness which can affect axon pathfinding [38-40]. The topography of the environment has been also identified as a potent guidance cue for growing axons and migrating cells through a process called contact guidance [41,42]. To dissect out the role of such physical cues, several engineered topographical substrates, such as grooves [43,44], pillars [45,46,47], pits [48,49] or synthetic fibers [50,51], were used to demonstrate *in vitro* the influence of topography on the migration of different cell types. A few studies employed micro-channels as a culture system for migrating cortical interneurons [52,53]. However, none addressed the potential influence of topographical cues in the migration of these cells.

In this study, we investigated this question using microstructured substrates and demonstrated the high sensitivity of cortical interneurons to the architecture of their environment. Our results revealed that topographical constraints efficiently modulate the migratory behavior of interneurons by controlling cell morphology, branching dynamics and cytoskeletal organization. In particular, by using two shapes of micron-sized pillars, we have been able to select and stabilize *in vitro* two main migratory behaviors observed *in vivo*, associated with specific morphologies: non-branched cells with long leading processes showed a directional movement among square pillars, and branched cells with short leading processes displayed a fast, exploratory migration among round pillars. This study therefore provides new insights into the guidance and the biology of these cells. The *in vitro* model described in this study thus emerges as a new promising tool to control and study interneuron migratory behaviors in physiological or pathological conditions.

## MATERIALS AND METHODS

### Animals

E12.5 pregnant Swiss mice were purchased from Janvier Labs. Animal experiments followed the French and European regulations for the care and protection of the Laboratory animals (EC Directive 2010/63, French Law 2013-118, 6 February 2013), and were authorized by the Ethical Committee Charles Darwin N◦5 (E2CD5).

### Microfabrication

The original master mold was fabricated by photolithography as follows. A 10μm-thick layer of SU8-2010 (MicroChem, USA) was formed on a 2-inch silicon wafer (Neyco, France) by spincoating at 3000 rpm for 30 s. After a soft bake (1 min at 65°C and 2 min at 95°C), the wafer was exposed to UV light (MJB4 Mask Aligner, 23mW/cm2 power lamp, SUSS MicroTec, Germany) through a hard chromium mask prior to a post exposure bake (1 Min at 65°C and 3 min at 95°C) and developed 2.5 Min under agitation using SU-8 developer (MicroChem, USA). After a 30 second oxygen plasma treatment, the obtained mold was exposed to a vapor of (Tridecafluoro-1,1,2,2 tetrahydrooctyl) trichlorosilane (AB111444, ABCR GmbH & Co. KG, Germany) for 20 minutes once, and used repeatedly without further treatment. The same procedure was applied once on every PDMS replica subsequently fabricated before another round of molding.

Polydimethylsiloxane (PDMS Sylgard 184, Sigma Aldrich) was mixed with its linker (ratio 1:10), poured onto the original master mold, degassed under vacuum and reticulated overnight at 70°C. Another round of PDMS molding and demolding was performed under the same conditions to get a final master mold with a negative topography. To create the final culture coverslip, PDMS was mixed with its linker, directly poured on the mold and spin coated at 3000 rpm for 30 seconds (for a final theoretical thickness of 30μm). Before reticulation overnight at 70°C, a glass coverslip was gently placed on top of the PDMS layer (See Fig.1A). After reticulation, the glass coverslip attached to the microstructured PDMS layer was gently demolded with a scalpel and isopropanol to facilitate detachment. Microstructured coverslips were then sonicated for 10 minutes in ethanol for cleaning and rinsed in water.

### Culture of MGE explants

Microstructured coverslips were fixed with dental silicon at the bottom of perforated petri dishes, which were used as a culture chamber for the video microscopy experiments. Otherwise, coverslips were placed in 4-well boxes.

#### Coverslip coating

N-Cadherin and laminin coating was performed according to the protocol detailed in [54]. Briefly, microstructured coverslips were first incubated for 6 hours at 37°C with 200μg/mL of poly-L-lysine (MW 538 000, Sigma). After 3 washes with water and drying for 15 minutes, coverslips were incubated overnight at 4°C with 4μg/mL laminin (from mouse sarcoma, Sigma) and 4μg/mL goat anti-human Fc antibody (Sigma) in borate buffer. Coverslips were then incubated with 0.25mg/cm2 purified N-Cadherin-hFc chimera protein (R&D system) for 2h at 37°C. After one wash with borate buffer, the coverslips were incubated at room temperature with a saturation buffer composed of 1.5% Bovine Serum Albumin (BSA) in borate buffer for 10 minutes and directly replaced by culture medium (all products from Gibco): Dulbecco’s Modified Eagle Medium Nutrient Mixture F-12 (DMEM/F12) supplemented with GlutaMAX supplement, HEPES buffer 10mM, glucose 30%, penicillin/streptomycin, N2 and B27 supplements.

#### Primary cultures

Medial ganglionic eminences (MGE) were extracted from the brain of E12.5 wild type embryos, and cut in 6-8 pieces of equal size (each one containing a part of the ventricular zone) as explained in [54]. All the explants from one MGE were placed on the surface of a microstructured coverslip at the bottom of the culture dish. After 6-8 hours in culture at 37°C, the first interneurons started migrating out of the explants.

### Time-lapse recording

Time-lapse recording started after 6-8 hours of culture, when the first migrating interneurons were seen exiting the explants. Before time-lapse recording, medium was replaced by a culture medium without phenol red, with sodium pyruvate 100mM (Gibco) and with an increased concentration of Hepes buffer (20 mM instead of 10 mM). Cultures were imaged in phase contrast on an inverted video-microscope (DMIRE2, Leica Microsystems) equipped with a Cool snap Myo camera (Photometrics, USA). Cells were recorded using a 20X objective (Leica, NA=0.7), every 2 minutes, for 12h. Acquisitions were controlled using the Metamorph software (Molecular Devices, USA).

### Immunostaining

For immunostaining, explants cultured for 24h were fixed in warmed (37°C) 4% paraformaldehyde (PFA), 1% sucrose in 0.12M phosphate buffer at pH7.4. After 5h in PFA, coverslips were rinsed 3 times in Phosphate-Buffered Saline 1X (PBS), and gently detached from the culture dish and put in a humid chamber. Cultures were incubated for 1h in a blocking solution containing 0.25% Triton, 10% normal horse serum (HS), 2% bovine serum albumin (BSA) in PBS. Cultures were then incubated overnight at 4°C with primary antibodies diluted in a solution containing 0.25% Triton, 2% HS, 1% BSA. The following primary antibodies were used: rat anti-tyrosinated tubulin (YL 1/2 ab6160, Abcam) 1:2000, rabbit anti-detyrosinated tubulin (L15, generous gift of Annie Andrieux, GIN, France) 1:1000. Primary antibodies were detected using anti-rat and anti-rabbit Alexa Fluor 488- or Cy3-conjugated secondary antibodies (Jackson Immunoesearch) diluted 1:1000 in PBS Triton 0.25% and incubated 1h30 at room temperature. Bisbenzimide (1:5000 in PBS Triton 0.25%, applied 5 minutes at room temperature) was used for fluorescent nuclear counterstaining. Cultures were mounted in Mowiol-Dabco (Sigma) and observed on an upright fluorescent microscope (DM6000; Leica Microsystems) equipped with 10x (NA=0.3), 20× (NA = 0.7), 63x (NA=1.4) and 100x (NA=1.4) objectives and a Coolsnap EZ CCD camera (Photometrics, USA) controlled by the Metamorph software.

### Scanning electron microscopy

6

Cultures were fixed in 2% glutaraldehyde in 0.1M phosphate buffer at pH 7.4. They were dehydrated in a graded series of ethanol solutions, then dried by the CO2 critical-point method using EM CPD300 (Leica Microsystems). Samples were mounted on an aluminum stub with a silver lacquer and sputtercoated with a 5nm platinum layer using EM ACE600 (Leica Microsystems). Acquisitions were performed using a GeminiSEM 500 (Zeiss).

### Structured-illumination microscopy (SIM)

For SIM, cultures performed on N-cadherin and laminin coated glass coverslips were fixed with PFA, and immunostained as described before. Super-resolution structured-illumination microscopy was performed on an Elyra PS.1 system (Zeiss) equipped with 63x (N.A. 1.4) and 100x (N.A 1.46) objectives and an iXon 885 EMCCD camera (Andor, Oxford Instruments).

### Data analysis

#### Fixed cultures of MGE cells

Quantifications were performed on images acquired with a 20X objective, either on mosaic pictures of whole explants or at the front of migration, in the 200μm-wide outermost ring around the explant. Explants from at least 3 independent experiments were analyzed in each condition.

To determine the mean distance of migration, the distance from the periphery of the explant to the most distant migratory cell was measured on 4 orthogonal directions in each explant and averaged. To calculate the internuclear distance, the coordinates of the cell bodies positions were extracted using ImageJ software. A custom-written MATLAB (MathWorks) script was used to calculate the distances between cell bodies and displayed for each cell the distance to its five closest neighbors. The cell orientation was measured after rotating images to have the same orientation of the pillars. Leading processes were manually traced and their orientations (expressed as angles) calculated using ImageJ (see Fig. 2A for definition). For multipolar cells, the main leading process was defined as the active process capped by the largest growth cone. Negative angles were converted to their symmetric positive counterpart to work in a half circle of 0-180° angles. Polar histograms were generated in MATLAB.

Leading and trailing process lengths, leading process width, as well as the number of branches and bifurcations of leading processes were measured and counted manually with the NeuronJ plugin of ImageJ, using the criteria described above and in Fig. S3.

#### Living cell analyses

Dynamic analyses were performed in a sample of cells recorded in at least 3 independent experiments for each condition. Cells in the sample migrated forward at the start of the recording session with minimal contacts with other cells.

The tip of the leading process (LPT) was manually tracked on 2-3 hours sequences of movement using the MTrackJ plugin of Image J. For unipolar cells, the LPT corresponds to the base of the unique growth cone (see scheme in Fig.6). For branched cells, the LPT was retroactively tracked as the base of the main growth cone capping the stabilized and selected branch. The directionality ratio (ratio between the distance from the first to the last point of the trajectory and the length of the actual cell trajectory) was calculated using the available Excel Macro in [55]. The quantification of LPT movement with respect to the pillars was performed using a custom-written MATLAB script. More precisely, a mask of the structures was extracted from the phase contrast movie, allowing to detect and count the pillars and to calculate the coordinates of their centroids. For each LPT position, the distance to the border of the closest pillar was measured on the line joining the LPT to the closest pillar centroid. The program displays a list of the LPT-pillar distances, as well as a graphical representation of the successive positions of the LPT in between the structures. On these graphics, the distance of the LPT to the closest pillar was color-coded

Growth cone divisions were surveyed for their time of appearance, their duration and their asymmetrical or symmetrical mode (see definition in Fig.4). From these data, the splitting frequency and the proportion of each splitting mode were calculated and averaged amongst the cells.

Cell body dynamics were analyzed in cells tracked for 6-8 hours. Cell tracking was performed manually using the MTrackJ plugin of ImageJ. Trajectories were color coded with the instantaneous migration speed using a custom-written MATLAB script. Directional reorientations and polarity reversals were counted manually and their frequency calculated for each cell. The directionality ratio was calculated using the Excel Macro in [55].

### Statistic

In text and figures, the variability around mean values is represented by s.e.m. (standard error of mean). Statistical analyses were performed using the GraphPad Prism software. Student’s t-test was used to compare null hypotheses between two groups for normally distributed data. Multiple groups were compared by ANOVA, followed by the Tukey’s posthoc test, or by the non-parametric Kruskal–Wallis test with Dunn’s posthoc test, according to the size and distribution of the data. Statistical differences between the frequencies of several parameters in distinct experimental conditions were assessed by a Chi2 test. Significance for individual values in distributions was assessed by the Fisher’s test. The number of data points for each experiment, the specific statistical tests and significance levels are noted in the legend of each figures.

## RESULTS

### Design of microstructured substrates to study the migration of interneurons

To test the role of topographical cues on cortical interneuron migration, we developed microstructured substrates of various topographies using photolithography and polydimethylsiloxane (PDMS) molding. The final culture coverslip is composed of a thin layer of microstructured PDMS strongly attached on a glass coverslip (Fig.1A). This original last step of fabrication allows high workability for cell culture because the glass/PDMS coverslips can be manipulated as regular glass coverslips. The 10μm high PDMS micron-sized structures provide a pseudo-3D open environment easily supplied with oxygen and culture medium. Fluorescent live or fixed cells can be imaged at various magnifications through the thin layer of PDMS, and immunostaining is easily performed. Before culture, the microstructured substrates were functionalized with a homogeneous coating of laminin and N-cadherin, two adhesion molecules promoting both interneuron polarization and migration [31,54,56]. Explants from the medial ganglionic eminence (MGE), the main source of cortical interneurons, were dissected from mouse brain embryos. These explants released polarized and radially-oriented interneurons that colonized the surrounding substrate for approximately 24 hours (Fig.1B). Regardless the topographical feature, the height of the structures (10 μm) constrained cell adhesion and migration at the bottom and on the edge of the microtopographies. The space in between structures defined the path available for cell migration.

We designed and tested different geometries of topography inspired from the literature, using flat PDMS surfaces as a control (Fig.1C and Fig.S1A). Grooves have been widely employed to promote the migration and guidance of different cell types [41]. We therefore first designed and tested a topography of grooves (6μm wide,10μm high) radiating from the explant and intersected every 200μm by concentric circular grooves, allowing possible cell reorientations as observed *in vivo* (Fig.1D and Fig.S1B). On this pattern, interneurons most often appeared as bipolar cells with long processes migrating in chain inside the grooves. A few isolated cells were observed at the front of migration (Fig. S1B). Some interneurons turned at intersections while their leading process remained non-branched, a case rarely observed *in vivo*. Because interneurons did not present their physiological mode of branching on this first pattern, we therefore decided in a second design to both increase the width and overall surface of migration paths and to increase the frequency of turning angles. This was achieved by implementing an array of hexagons (68μm long, 10μm high) providing 10μm wide intersecting paths forming 100-degree open angles (Fig.1E and Fig.S1C). We observed that the leading processes of interneurons now branched at bifurcations. However, similarly to what was observed on flat surfaces, cell bodies often aggregated at the front of migration, preventing a correct visualization of individual cells.

**Figure 1:**
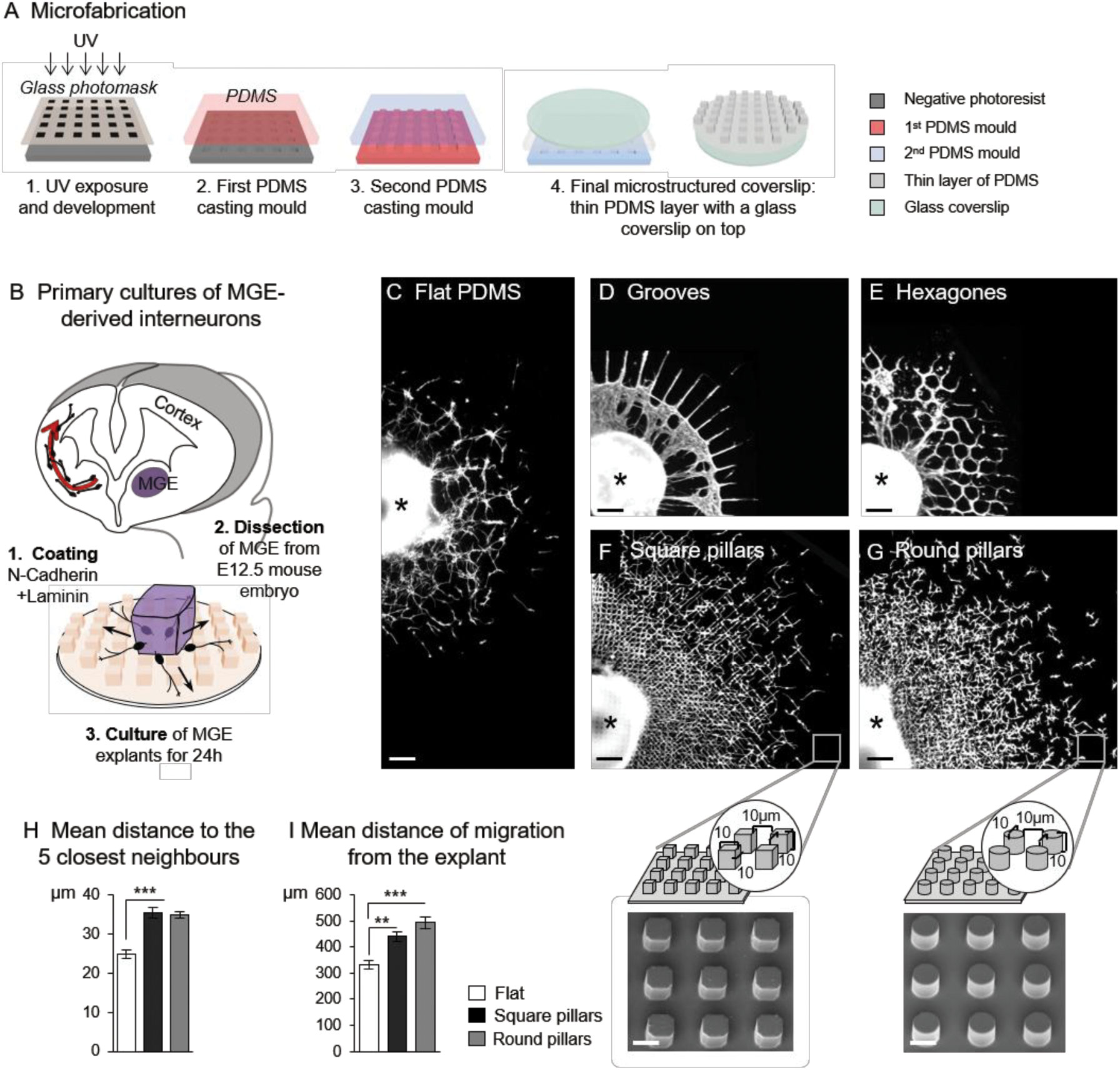
In vitro migration of interneurons on microstructured substrates. (A) Schematics of the microfabrication steps used to create the microstructured coverslips. (B) Principal steps of the primary culture of MGE-derived embryonic interneurons. (C-G) Distribution of migratory interneurons on flat PDMS (C), in 6μm wide grooves (D) in between hexagons (E), and in between square (F) and round pillars (G) with schematics and scanning electron microscopy (SEM) images of the pillared microstructures. Pictures in (D-G) illustrate the upper right quadrant of cultures of MGE explants (black star) fixed after 24h *in vitro* and immunostained with antibodies against tyrosinated tubulin. Scale bars, 100μm (C-G), 10μm (SEM pictures). (H) Mean distance of each cell to its 5 closest neighbors at the front of migration on flat and pillared substrates. Measures were performed around 10 explants in 4 (flat) or 3 (pillars) independent cultures. Error bars represent mean ± SEM. One-way ANOVA, Tukey’s post test (***, p<0.0001). (I) Mean distances measured as explained in Methods were averaged for 49 explants (4 cultures) on flat PDMS, 98 explants (7 cultures) on square pillars, 62 explants (6 cultures) on round pillars. Error bars represent mean ± SEM. Non-parametric ANOVA, Dunn’s post test (*** p<0.0001, ** p<0.001).

We hypothesized that such aggregation phenomena may arise from the confinement induced by the hexagonal grooves and the still too restricted migration surface they provide. We therefore replaced grooves with features of smaller size while keeping the same inter-topography spacing. This third design consisted in regularly spaced micro-pillars, square or round, of 10μm in width/diameter and height, separated by 10μm (Fig. 1F,G). In these microenvironments, interneurons leaving MGE explants migrated in between pillars showing more physiological branched morphologies (Fig.S1D). Cells no longer aggregated and remained individualized at the front of migration. Accordingly, the mean distance between neighboring cell bodies at the front of migration on pillars was significantly increased as compared with cultures performed on a flat substrate (Fig.1H). In addition, the distance of migration from the explant after 24 hours was also significantly higher in both types of pillars as compared with the flat surfaces (440 ± 18μm and 493 ± 21μm in square and round pillars respectively vs 333 ± 16μm on flat, Fig.1I).

In conclusion, the use of regular arrays of micropillars improved the overall migration of interneurons on adhesive substrates, promoting a physiological-like migratory behavior of individual cells. We therefore used these pillared substrates to perform an in-depth analysis of the influence of the topography on the migratory behavior of embryonic interneurons.

### Microtopography restricts interneuron orientation and morphology

Interneurons migrating away from MGE explants showed various macroscopic organizations. Interestingly, interneurons migrating on square pillars frequently formed a mesh of long and perpendicularly oriented processes. In contrast, the majority of MGE explants placed on round pillars released interneurons with shorter processes more randomly oriented (Fig.S2).

We therefore analyzed the orientation of the main leading process of interneurons located at the front of migration using fixed cultures performed on microstructured and flat surfaces (Fig.2A). In cultures performed on a flat PDMS surface, interneurons were randomly oriented. Of note is that pillared substrates displayed four lines of symmetry (i.e. horizontal, vertical and diagonals). Interneurons in between round pillars oriented equally along the four directions corresponding to the aforementioned lines of symmetry (Fig. 2B). Remarkably, when round pillars were replaced by square pillars, the orientation of interneurons was almost exclusively restricted to the two diagonals (Fig.2B, with 80% of the cells at the front of migration oriented in the {30-60°} and {120-150°} ranges). The topography of the substrate and the shape of the structures have therefore a major impact on the orientation of the interneuron leading process. To get a better insight into these remarkable differences, we next analyzed interneuron morphology on the different substrates.

**Figure 2:**
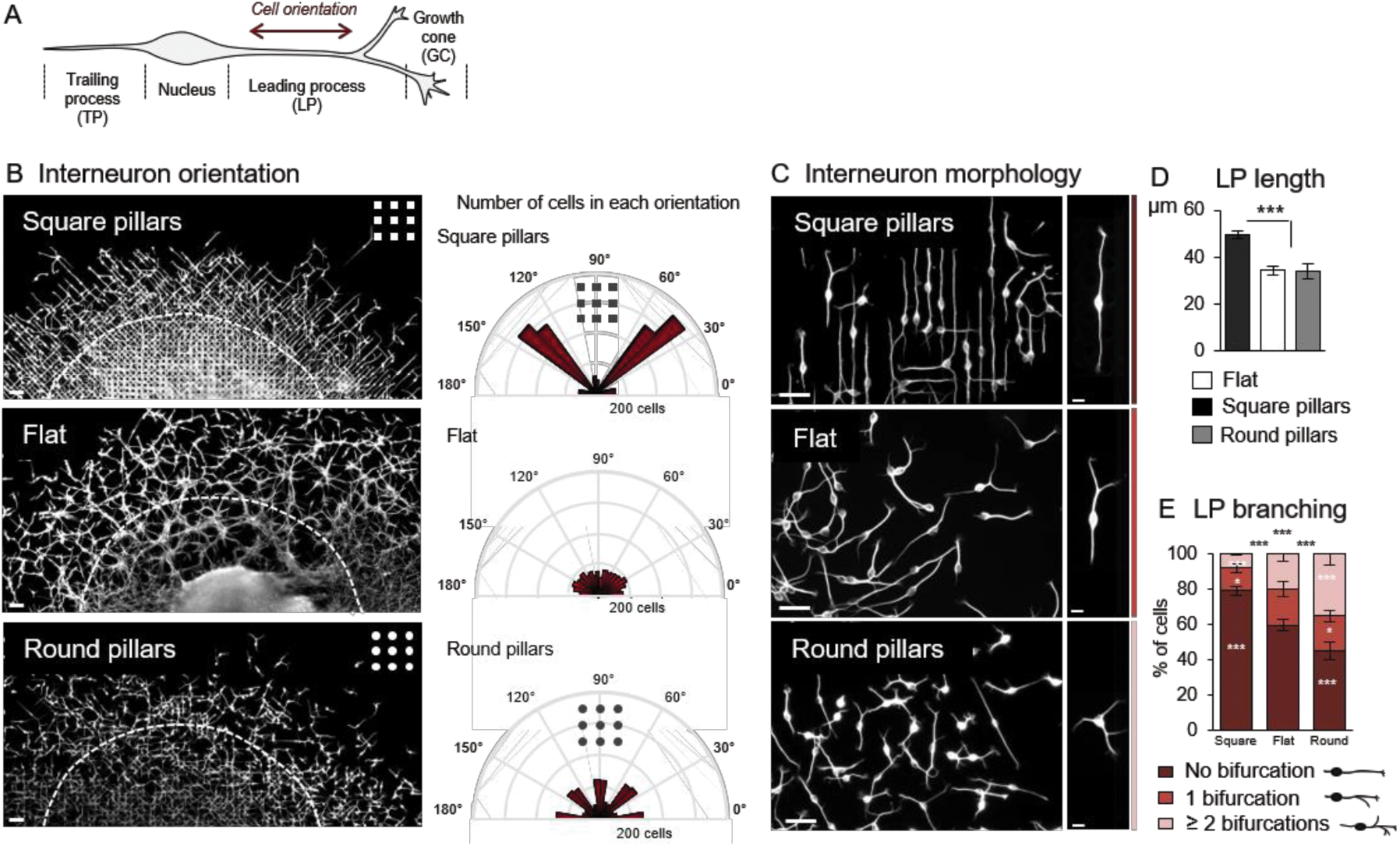
Microtopography influences interneuron orientation and morphology. (A) Schematics of the typical morphology of a migratory interneuron. Interneuron orientation is defined as the orientation of the proximal part of the leading process. (B) Left: partial view at low magnification of interneurons surrounding an explant cultured for 24h on either square pillars (top), flat surface (middle), or round pillars (bottom). Interneuron orientation was measured in the peripheral area limited by the dashed white line (3200 cells per condition;Flat: 5 cultures, square pillars: 3 cultures, round pillars: 4 cultures). Right: Polar plots show the number of cells in each class of orientation. The distribution of cell orientations on square pillars significantly differs from the distributions on flat surface or round pillars (Chi2 test, p<0.0001). (C) Pictures show the morphologies of interneuron at their front of migration on pillared and flat surfaces. (D) Average length of the leading process (from cell body front to the base of the largest growth cone) measured at the front of migration in 3 independent cultures (square pillars, 411 cells; flat PDMS, 436 cells; round pillars, 379 cells). Error bars represent mean ± SEM. Non-parametric ANOVA, Dunn’s post test (***p<0.0001). (E) Percentage of 3 main classes of cell morphologies defined by the number of bifurcations of the leading process, on each substrate of migration. Analyses were performed in 3 independent cultures (Square pillars, 411 cells; flat PDMS, 436 cells; round pillars, 379 cells). Error bars represent mean–SEM. Couples of distributions were compared using a Chi2 test (black stars, *** p<0.0001). White stars within columns compare cells from the same morphological type on the flat PDMS (Fisher test, *** p<0.0001, *p<0.05). Interneurons were immunostained with antibodies against tyrosinated tubulin. Scale bars, 50μm (B), 30μm (C, left panels), 5μm (C, right panels)

The leading processes of interneurons cultured on square pillars were significantly longer on average (49.6±1.5μm) than those of interneurons cultured on either flat surfaces or round pillars (34.2±1.7μm and 33.7±3.3μm, respectively, Fig 2C,D). In addition, the proportion of cells with a single non-bifurcated leading process was higher on square pillars (79% ± 2.8) than on both the flat surfaces and round pillars (59.7% ± 3.35 and 45% ± 4.86 respectively, Fig.2D). Conversely, the proportion of branched interneurons showing at least 2 bifurcations of their leading process was significantly increased on round pillars (35% ± 6 on round vs 8% ± 0.5 on square pillars, Fig.2E). On a flat surface, the proportion of these different morphologies was intermediate (Fig.2E). The morphological diversity observed on flat surfaces appeared thus strongly reduced on pillared surfaces. Strikingly, non-branched cells with a long leading processes represented the large majority of migratory cells on square pillars whereas short and branched cells were preferentially found on round pillars (Fig.2C, Fig S3D).

Analyses of morphological features associated with the leading process length confirmed that interneurons with long leading processes had fewer lateral branches. These neurons display in addition a thinner leading process, a more elongated nucleus and a longer trailing process than interneurons exhibiting short leading processes (Fig.S3E-H). Interestingly, interneurons migrating on a flat surface can successively adopt a long non-branched and a short and branched morphology (Fig.S3A). Because cultures on square and round pillars were respectively enriched in long non-branched and short and branched interneurons, we concluded that square and round pillars each stabilized opposite morphologies of interneurons that likely represent the two extremes of a morphological spectrum exhibited by a unique population of migrating interneurons.

### Interneuron positioning within pillared surfaces

We then examined in detail the positioning of interneurons with respect to the microtopography of pillars using scanning electron microscopy (SEM, Fig.3). SEM observations confirmed that long cells migrating among square pillars had an elongated soma that often contacted simultaneously two pillars positioned on each side of the cell (Fig.3A), contrasting with the spherical soma of the branched cells migrating among round pillars that predominantly engaged with a single pillar at a time (Fig.3B). These observations were further confirmed by dynamic studies (Fig.S4).

**Figure 3:**
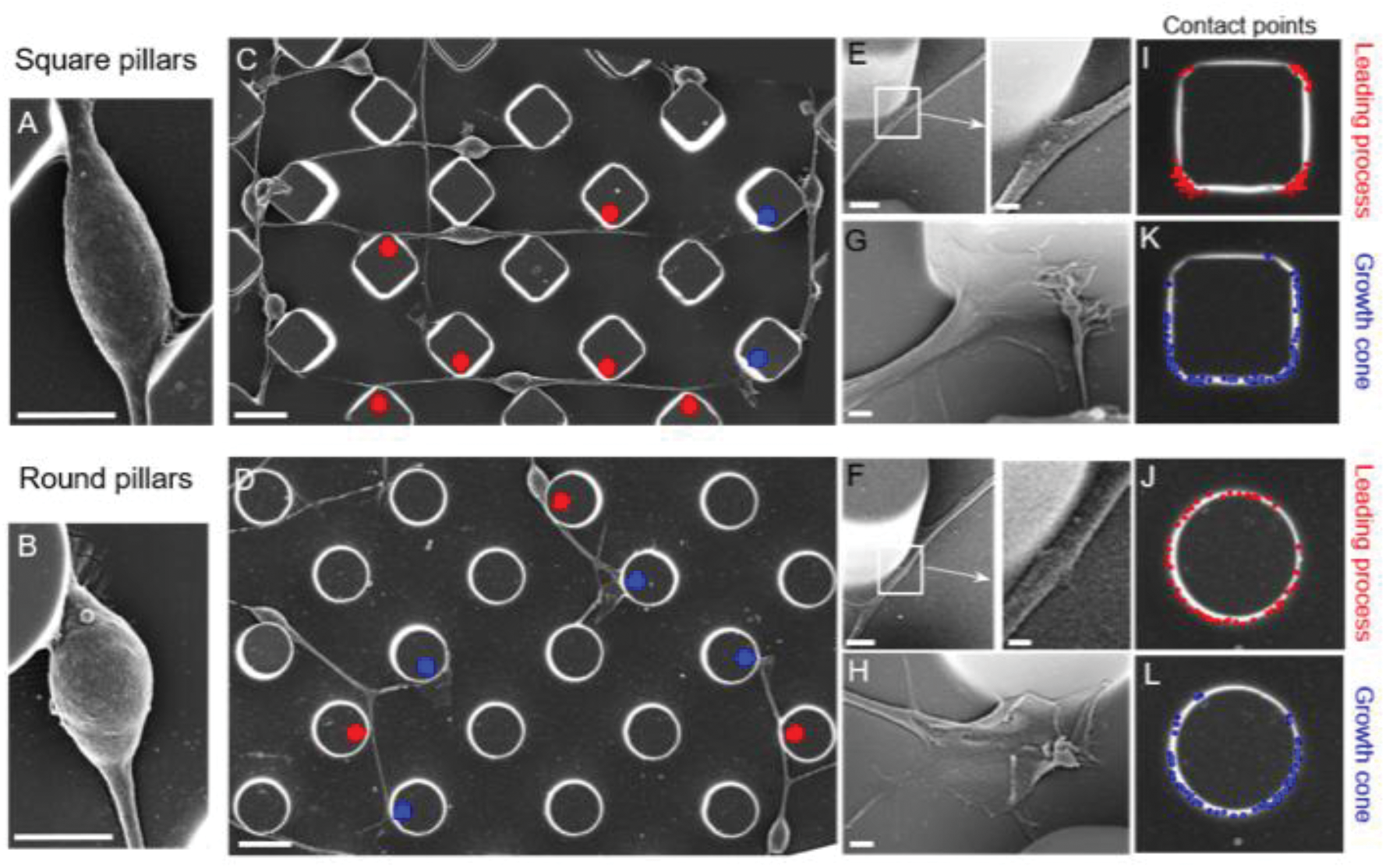
Scanning electron microscopy (SEM) analysis of interneuron positioning among square (A,C,E,G) and round (B,D,F,H) pillars. Representative pictures of contacts between pillars and the cell bodies (A,B), and the neuritic processes (CL) of interneurons. In (C,D), red dots show the contact points with the leading processes and blue dots, the contact points with the growth cones. (E,F) show leading processes interaction with the pillars and (I,J) their points of contact with the structures. (G,H) show growth cones morphologies on both pillared surfaces and (K,L) the contact points of growth cones with the structures. The contact points of leading processes and growth cones with the pillars were plotted for 60 cells per condition in 3 different experiments. Scale bars, 4 μm (A,B), 10μm (C,D), 2μm (E,F), 0.5μm (E,F, enlarged views), 1μm (G,H).

The leading processes stereotypically extended from corner to corner of the square pillars (Fig.3C, red dots), as shown by the restricted contacts of the leading process with the corners of the structures (Fig. 3E,I). At these corners, processes presented small enlargements suggestive of specific interactions with the structures in this region. On the contrary, leading processes extending at the basis of round pillars contacted the pillars on their whole circumference, surrounding them (Fig. 3D,F,J). Growth cones, the protrusive and exploratory structures at the extremity of the leading processes, showed very complex and diverse shapes in both types of microtopographies (Fig. 3G,H). They often positioned along the vertical side of pillars and could largely enwrap them regardless of their round or square shapes. The contact points of growth cones were regularly distributed all around the round and square pillars, suggesting a wide and random exploratory behavior of the growth cones (Fig.3K,L) that contrasted with the restricted interaction of the leading processes with the corners of the square pillars.

### Microtopography influences growth cone divisions

As different branching modes were observed on round versus square micropillar shapes, we examined the dynamics of this major morphological trait in interneurons using time-lapse sequences of 2-3 hours. A careful examination of leading growth cone divisions revealed two main modes of splitting, either symmetrical that produced sister branches with similar activities (pink arrows in Fig.4A, see Movie1) or asymmetrical that gave rise to a large growth cone and a much thinner and transient lateral process (Fig. 4B, see Movie 2). While these two splitting modes were present in similar proportions on flat surfaces (symmetric: 47.7% ± 4.7, asymmetric: 40.7% ± 4.5, Fig.4C), symmetric splitting was dominant in round pillars (symmetric: 56.1%±3.9, asymmetric: 26.7% ± 4.2). On the contrary, asymmetric splitting was preferentially observed on square pillars (symmetric: 30.6% ± 6.1, asymmetric: 51.1% ± 6.1). In addition, growth cone divisions were twice less frequent on square pillars than on flat surfaces and round pillars (1.42 bifurcation/h ± 0.23 in square, 2.44 ± 0.30 on flat, 2.77 ± 0.30 in round, Fig. 4D). We moreover noticed that asymmetric divisions often associated with a compact morphology of the growth cone. Accordingly, tyrosinated microtubules invading the growth cone of interneurons migrating between square pillars spread less largely than in the growth cone of interneurons migrating between round pillars (Fig. 4E).

**Figure 4:**
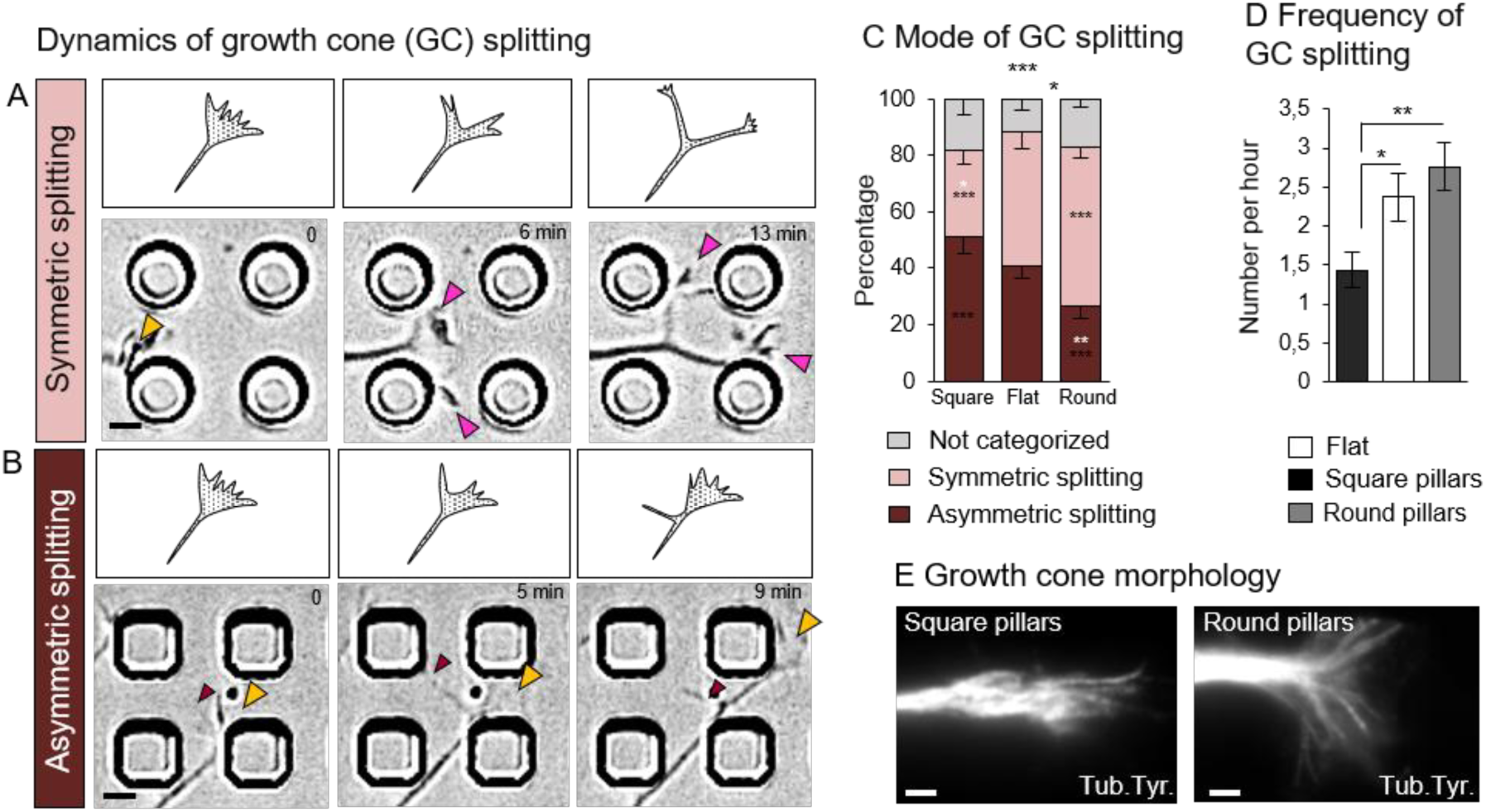
Microtopography influences growth cone splitting and branch formation. In (A,B), schemes (top) and time lapse sequences (bottom) illustrate two preponderant modes of growth cone splitting and branch production exhibited by migrating interneurons. A: symmetric growth cone splitting produces two active growth cones (pink arrows) that generate two symmetrical branches. Yellow arrows indicate a main growth cone. (See Movie 1). B: asymmetric growth cone splitting produces a lateral filopodium (brown arrow) that becomes a thin lateral branch quickly eliminated. (See Movie 2). (C,D) Histograms report the percentage of asymmetric and symmetric growth cone splitting (C) and the frequency of growth cone splitting (D) observed during recording sessions of 2-3 hours in cultures performed on the square pillars (23 cells, 6 cultures), flat PDMS (23 cells, 4 cultures), and in the round pillars (30 cells, 5 cultures). C: error bars represent mean–SEM. Splitting modes significantly differ between both pillared substrates (Chi2 test, black stars above columns, ***p=0.0002, * p=0.02). Stars within columns indicate differences within a single type of splitting mode (Fisher test). Black stars compare with the other type of pillars, white stars compare with flat (*** p<0.0004, ** p=0.0073, * p<0.03). D: error bars represent mean ± SEM. Branching frequencies were compared using a nonparametric ANOVA, Dunn’s post test (p=0.0019). (E) Tyrosinated tubulin immunostaining of growth cones showing representative morphologies. Scale bar, 2μm.

In conclusion, the differences in the mode of growth cone splitting observed in both pillared surfaces support the establishment of differently branched morphologies at the population level.

### Microtopography influences interneuron dynamic behavior

The global migratory behavior of interneurons was analyzed by phase contrast video-microscopy on sequences of 6-7 hours movement by following individual cells at the front of migration and analyzing the cell directionality and motility parameters. Typical behaviors observed in both pillared and on flat surfaces are displayed in Fig. 5A-C, representing the color coded cell body trajectories with the instantaneous speed, and in Fig. 5D-F where short temporal sequences of interneuron movements on pillared and flat surfaces are reproduced.

**Figure 5:**
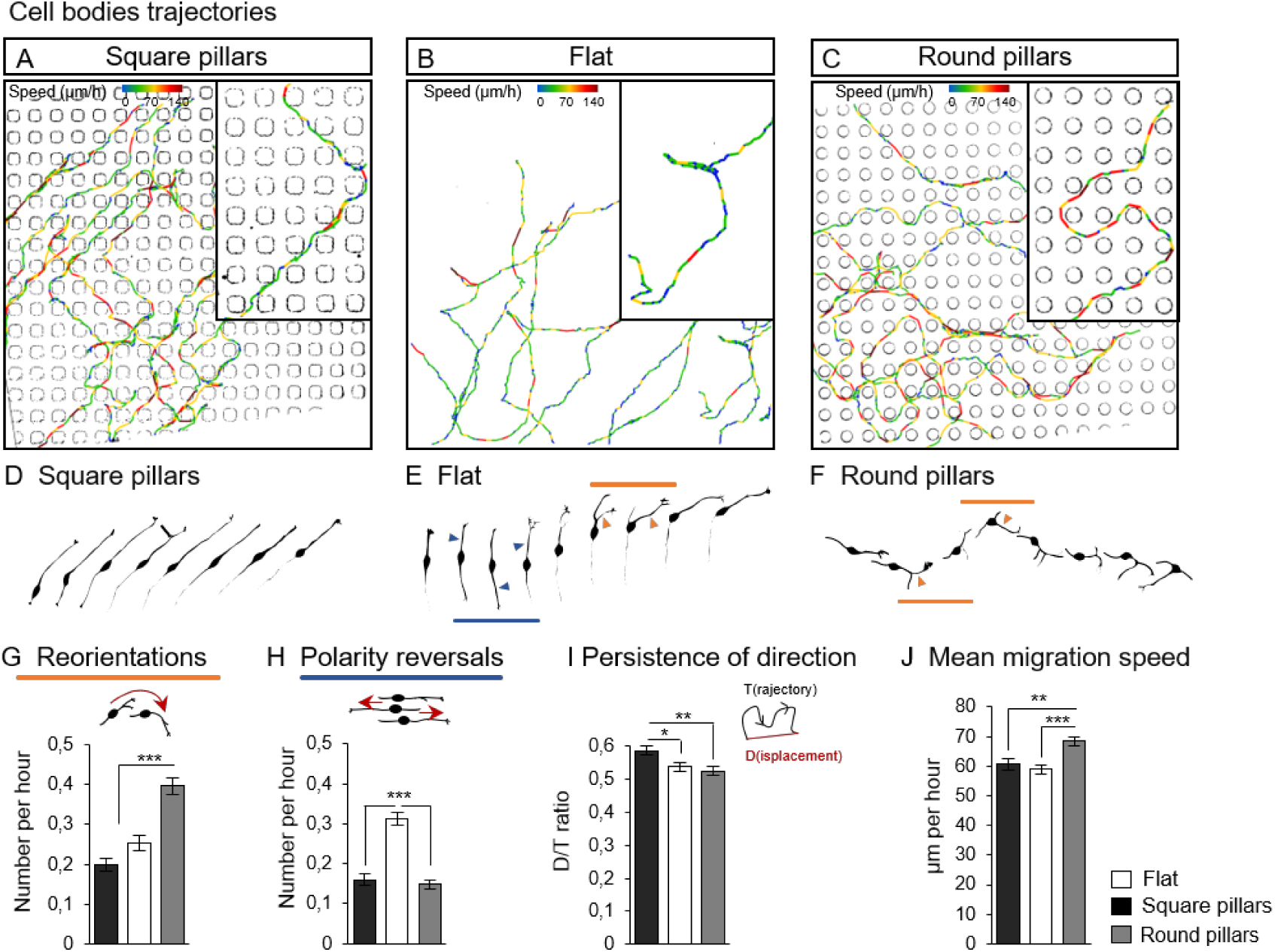
Microtopographies influence interneuron dynamics. (A-C) Representative examples of cell body trajectories in square (A) and round (C) micropillars, and on flat PDMS (B). (Representative examples in Movies 3 (flat), 4 (square pillars), and 5 (round pillars)). Trajectories are color coded for the instantaneous migration speed of the cell body. (D-F) Drawing of time lapse sequences representative of interneuron migratory behaviors on each substrate. Time interval, 18 minutes. Blue arrows indicate polarity reversals; orange arrows indicate directional changes operated by the selection of a diverging branch. (G,H) Mean frequencies of reorientations (G) and of polarity reversals (H) were compared on square pillars (118 cells, 7 cultures), flat PDMS (114 cells, 5 cultures), or round pillars (n=142 cells, 6 cultures). Significance of the results was assessed using a non-parametric ANOVA followed by a Dunn’s post test (p<0.0001). (I) Mean persistence of direction expressed as the ratio between the cell displacement D and the length of the actual trajectory T. Non-parametric ANOVA and Dunn’s post test (p=0.0063). (J) Differences in mean instantaneous migration speeds were assessed using a non-parametric ANOVA and Dunn’s post test (p=0.0002). In histograms, error bars represent the mean ± SEM.

On flat surfaces (Fig. 5B, E and Movie 3), interneurons alternated between directional phases (when the soma moved forward in a non-branched leading process, sequence in-between blue and orange arrows in Fig.5E) and phases of reorientations (when a lateral divergent branch was selected for soma translocation, orange arrows in Fig. 5E). Changes in direction could also arise from a polarity reversal between leading and trailing processes (blue arrows in Fig. 5E). In square pillars, interneurons presented long directional phases along a diagonal, interrupted by a few turns (often at 90°, Fig. 5A). Interneurons showed infrequent phases of reorientations, in agreement with their non-branched morphology (Fig.5D, G and Movie 4). In contrast, interneurons in between round pillars, which were often branched, showed accordingly an increased frequency of reorientations and erratic trajectories with directional phases of short duration (Fig. 5C, orange arrows in Fig.5F and Movie 5; 0.39 reorientations/h ± 0.02 vs 0.25 ± 0.01 on flat and 0.19 ± 0.01 on square, Fig.5G). Interestingly, the frequency of polarity reversals was very low on both microstructured substrates as compared to the one on the flat surfaces (0.16 ± 0.01 and 0.15 ± 0.01 reversals/h in square and round respectively vs 0.3 ± 0.01 on flat, Fig.5H), showing a positive effect of micropillars on cell polarity stabilization. The persistence of direction (measured by the directionality ratio, Fig. 5I) was higher in square pillars (0.59 ± 0.01) than in round pillars (0.52 ± 0.01) or on flat surfaces (0.53 ± 0.01). Finally, we analyzed the cell motility and found that the mean migration speed of cells within square pillars and on flat surfaces did not differ significantly (60.68 ± 1.98 μm/h in square pillars, 58.9 ± 1.6 μm/h on flat surfaces), whereas cells migrating within round pillars moved significantly faster (68.4 ± 1.54 μm/h, Fig.5J). On the three types of substrates, cell soma alternated between fast and slower migration movements, a cardinal property of the dynamics of interneuron somas (see color coded trajectories in Fig. 5A-C).

Overall, the two morphological types of migratory interneurons enriched in both micropillared environments associate with two distinct migratory behaviors: slow and directional in between square pillars, faster but less directional in the round pillars. These results highlight a strong link between morphological and dynamic parameters.

To further decipher these different cellular behaviors, we next investigated the dynamic interactions of the cell tip with the microstructured environment.

### Cell tip navigation between pillars

Cell tip navigation around the structures was analyzed on sequences of movement recorded by phase contrast video-microscopy. Because the complex and fast transformations of the growth cone in migrating interneurons is incompatible with the resolution of recordings, our analysis of the cell tip is focused on the tracking of the growth cone base, corresponding to the leading process tip (LPT) (red dot on scheme in Fig.6 and Movies 6,7).

Trajectory analyses showed that despite occasional 90° turns, the LPT of interneurons migrating in-between square pillars exhibited straight trajectories and maintained the same direction of movement over several hours (Fig.6A). Accordingly, the persistence of direction measured by the directionality ratio was significantly higher on square pillars (0.80 ± 0.01) than on flat surfaces (0.64 ± 0.02, Fig. 6D). LPT movements among square pillars were often confined to the diagonal direction where they progressed from corner to corner (Fig. 6E,G, see Movie 6). Accordingly, the distance between a LPT and the closest pillar on its diagonal trajectory never exceeded 4 μm (mean amplitude 2.23 μm ± 0.09, Fig. 6I,K). Interestingly, this limited range of exploration of the LPT did not reflect the variety of growth cone positions observed by SEM around the square pillars (Fig. 3I). Rather, LPT trajectories prolonged the leading process orientation and we therefore conclude that LPT movements were highly constrained by the leading process behind.

**Figure 6:**
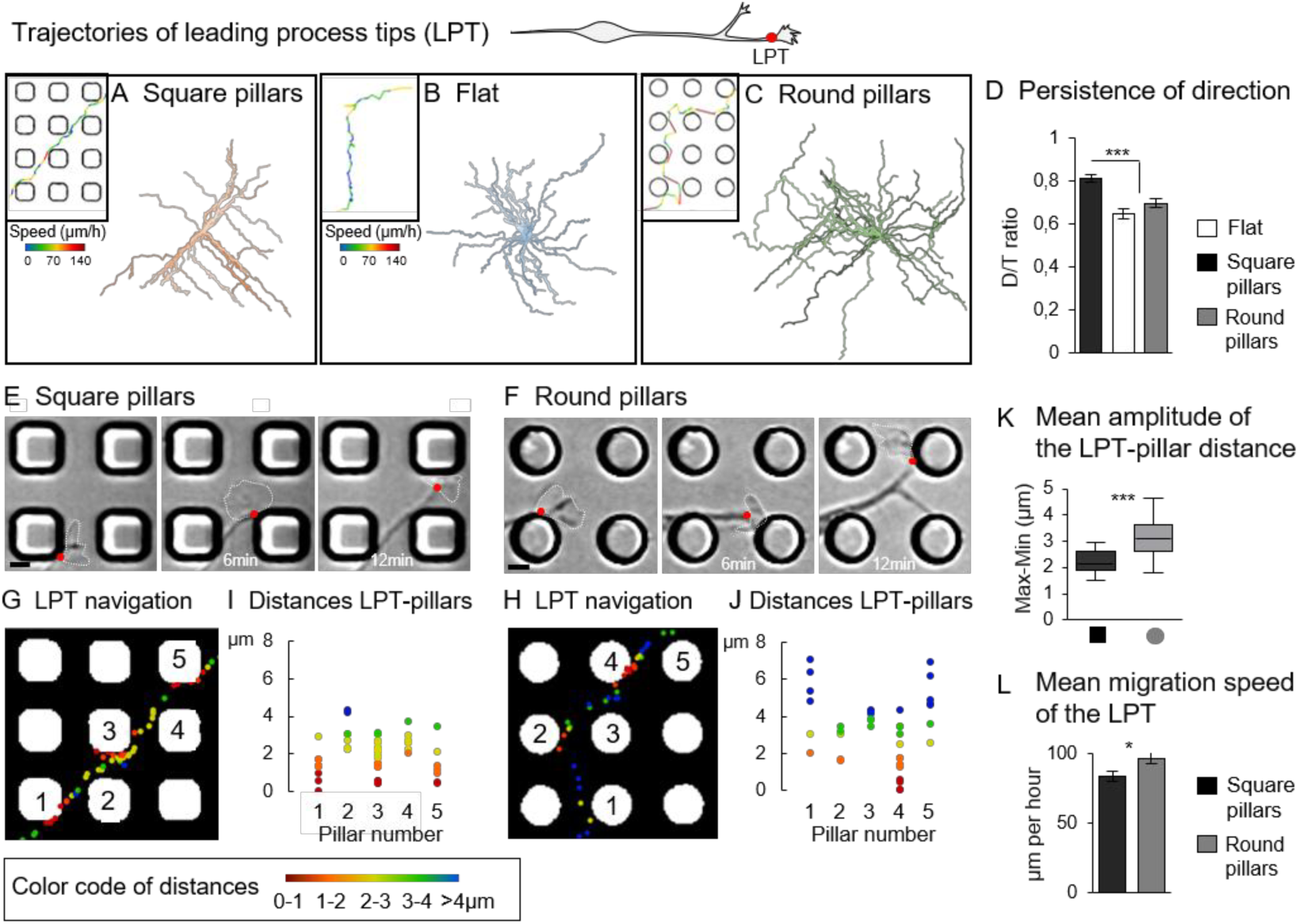
Cell tip navigation in between pillars. (A,B,C) Trajectories of the tip of the leading process (LPT on scheme) in between square pillars (A), on flat PDMS (B) and in between round pillars (C). Inserts show enlarged representative trajectories color coded with the instantaneous LPT migration speed. (D) Mean persistence of direction (see legend Fig.5I) of LPT migrating on square pillars (30 cells, 6 cultures), flat PDMS (23 cells, 4 cultures), round pillars (30 cells, 5 cultures). Error bars represent mean ± SEM. One-way ANOVA Tukey’s post test (p<0.0001). (E,F) Time lapse sequences illustrate representative LPT movements (red dots) among square (E) and round (F) pillars. See additional examples in Movies 6 and 7. Scale bars, 5μm. (G,H) Analysis of LPT navigation among pillars: the distance from a LPT to the closest pillar was calculated, color coded (see bottom scale), and the successive positions of the LPT represented with the color code. LPTs were recorded during 1hour 20 min, with a time interval of 2 minutes. (I,J) Plots show the dispersion of the LPT-pillars distances, which corresponds to the space explored around each pillar by the LPT. (K) The mean difference between the maximal and minimal LPT-pillar distance for each pillar, represents the amplitude of LPT exploration around pillars. Square pillars n=21 cells (6 cultures), round pillars n=29 cells (5 cultures). Student T-test (p<0.0001). (L) Histogram compares the migration speed of LPTs on round (30 cells, 5 cultures) and square pillars (29 cells, 6 cultures). T-test (p=0.0171). Error bars represent mean ± SEM

On the contrary, LPT navigating in-between round pillars explored the space between pillars in an apparent random way with frequent turns, as evidenced by their low persistence of direction (Fig. 6C,D, see Movie 7). The distances between the LPT and the closest pillar varied more largely in the round than in the square pillars (Fig 6F-K, mean amplitude: 3.076 μm ± 0.12). In addition, LPT migrated faster among round pillars than among square pillars (Fig.6L, 96.36 μm/h ± 4 vs 83.30 μm ± 3.47 in square pillars). The leading tips of interneurons migrating between round pillars were thus more motile and less directed than those of interneurons migrating between square pillars.

Overall, the dynamics of the leading tip appeared similar to the dynamics of the cell body in the same conditions (Fig.S4). However, the cell tip of interneurons migrating between square pillars displayed directional movements that seemed strongly constrained by the orientation and stability of the leading process behind.

### Microtubule cytoskeleton organization in interneurons on microstructured substrates

To gain insight into the structural organization of the leading process of interneurons migrating on square and round pillars, we next examined the organization of the microtubule network and tubulin post-translational modifications. Dynamic microtubules were immunostained with antibodies against tyrosinated tubulin. Antibodies against detyrosinated tubulin labeled stable microtubules (Fig7). Interneurons migrating in-between the square pillars presented leading processes enriched in long and tight bundles of stable microtubules (Fig. 7A) whereas interneurons migrating in-between round pillars showed wide leading processes enriched in dynamic microtubules organized in short and spread structures (Fig.7B). Interneurons migrating in both microtopographies thus differed by the relative abundance of stable and dynamic microtubules in their leading processes. Interestingly, the organization of the microtubule network at the level of the cell body also differed. In both cases, a network of looped stable (detyrosinated) microtubules surrounded the nucleus (Fig. S5A,B). This looped organization was further confirmed and detailed using SIM super resolution microscopy on a flat glass surface (Fig.S5C). In long cells migrating in-between square pillars, an alignment of dynamic (tyrosinated) microtubules positioned below the nucleus in contact with the substrate, forming a rail in apparent continuity with the microtubule bundles extending in the leading process and in the trailing process (Figs.7 A2 and S5,A, yellow arrows). Such an organization was never observed in the short-size branched cells (Fig.7 B2 and S5,B).

**Figure 7:**
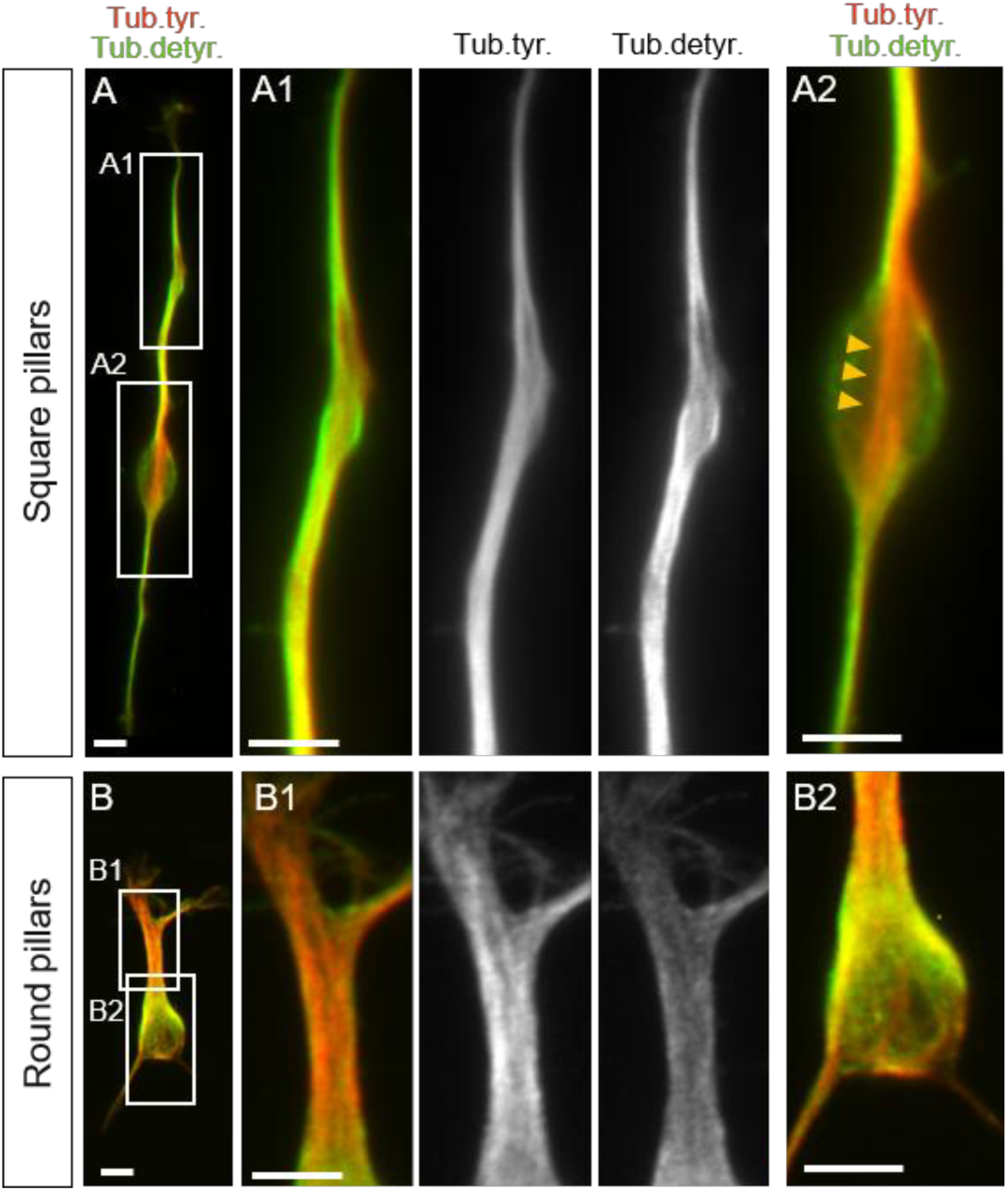
Microtubule cytoskeleton organization in interneurons on pillared substrates. (A,B) Migratory Interneurons on square (A) and round (B) pillars were immunostained with antibodies against tyrosinated tubulin (dynamic microtubules, red) and detyrosinated tubulin (stable microtubules, green). (A1,B1) are enlarged views of the leading process and (A2, B2) of the cell body. Panels show maximum projections of epifluorescence microscopy pictures acquired at 2 z levels. Scale bars, 5μm.

The navigation of a long and straight leading process with a specific and restricted interaction pattern with the corners of squared pillars appeared thus constrained by a network of stable and dynamic microtubules organized in continuous bundles from the cell body to the LPT. This property seems to provide the basis of the particular orientation and alignment observed in the diagonal direction of square pillars.

## DISCUSSION

The present study reveals the existence of a contact guidance phenomenon for embryonic cortical interneurons. Using microstructured substrates characterized by arrays of micron-sized pillars, we have shown that fundamental traits of the migratory behavior of embryonic interneurons are influenced in specific ways by the topography of the microenvironment. Beside the observation that pillared surfaces stabilized the polarity of interneurons, our most striking finding was that the directionality and dynamics of interneurons are sensitive to the detailed shape of the pillars. Remarkably, each pillared surfaces preferentially selected one out of the two opposite types of movements exhibited by migrating interneurons in physiological contexts e.g. directed and re-orienting movements [7,9,15,16]. Our approach therefore provides the usual advantages of *in vitro* platforms, such as accessibility, fine tuning of cell culture parameters and a precise quantification of various aspects of cell behaviors, but also allows the specific application of these tools to interneurons in different migratory stages.

Migrating interneurons extend a branched leading process. The directionality of their movements thus reflects their ability to select and stabilize a neuritic branch with a specific orientation, in which the nucleus translocates afterwards [57]. We showed that interneuron interactions with the square micropillars prevented growth cone divisions and thereby the production of divergent branches. In this environment, a unique long and thin leading process, in which the nucleus moved forward, elongated along the corners of the pillars. On the contrary, interactions with round pillars promoted growth cone divisions. Branches contacted any surrounding pillar and associated with fast forward movements of the nucleus. Square pillars thus favored directional forward movements of interneurons with a single and long leading process, whereas round pillars favored the exploratory movements of interneurons with several short-sized branches. Accordingly, live-cell recording in organotypic cortical slices has shown that the long-size and non-branched interneurons observed in the tangential migratory pathways efficiently progress in the tangential direction to invade the embryonic cortex, whereas interneurons with a shorter and branched leading process often operate directional change, for example switching from a tangential to a radial orientation near the cortical plate [7,9,10,58].

In the square pillared arrays, the long and thin leading processes contained bundled and stable (detyrosinated) microtubules. This particular organization associated with both asymmetrical splitting of the growth cone and decreased splitting frequency observed in this environment. In the round pillars, interneurons often presented a short, wide and branched leading process with the mildly bundled, dynamic microtubules associated with a rounded nucleus. This is consistent with the positive link between microtubule destabilization and branching reported in interneurons lacking Doublecortin (DCX), a protein associated with microtubules stabilization [59], and with directionality alterations in interneurons treated with increasing doses of nocodazole, a drug that depolymerizes microtubules [60]. Interestingly, microtubule detyrosination seemed to appear gradually from the periphery when the leading process length increased, suggesting that the stabilization of microtubules might be linked to the lengthening and thinning of the leading process.

The microtopographies therefore stabilized two different cellular states, which likely represent the two extremes of a wide intrinsic morphological and dynamic repertoire that a single population of interneurons can adopt. This raises the question of the selection of these morphological types by the different topographical shapes. The observation that the long leading processes extending in the diagonal direction of square pillars engage restricted physical contacts with the corners, in contrast with their ability to enfold round pillars, suggests that the curvature might be a key parameter. The localized high curvature at the corners of square pillars may prevent the bending of the leading process, a phenomenon already observed in axons [61], thus stabilizing its orientation and trajectory once engaged into a diagonal path. How regular and discrete contact points with the structures would in turn stabilize a long, stable and non-branched leading process remains however as an open question. The development of a mechanical tension in the leading process between the enlarged adhesive areas observed at the corners, in line with the elongated shape of the soma, is a hypothesis that would deserve further investigations. Interestingly, it has been shown that applying tension on growing axons by attachment to carbon nanotubes suppresses branching [62], similarly to what we observed in between square pillars.

Besides the interaction with individual microstructures, the global architecture of the pillar arrays must also be considered. In particular, we observed that surfaces patterned with square pillars induced physical constraints on the cell soma, which likely influenced their movements and motility. These constrains were mostly reduced within the round pillar arrays due to their reduced surface, providing thus an increase of free surface suitable for cell migration in-between pillars. This may facilitate the growth cone random exploration of the environment and the splitting of growth cones favoring multipolar cell morphologies and directional changes.

*In vivo*, interneurons alternate between globally directed movement (in the tangential pathways and radially within the cortical plate), and reorientations (towards the cortical plate from tangential paths). More generally, these different phases of movement can be seen as two different migration strategies: one directed and the other a more exploratory one, both serving different purposes along the course of interneuron displacement. The signals controlling the switch between directed movements and re-orientation remain partly elucidated [9,52,63-66]. Our study reveals that a simple change in the geometry of the topography can orient interneurons towards one strategy of movement or the other *in vitro*. *In vivo*, embryonic interneurons are exposed to various and constantly remodeled cytoarchitectural arrangements in the developing cortex. In light of our results, we can therefore hypothesize that the architecture of the environment can influence interneuron migration, and contribute to the selection of migratory behavior in the developing cortex.

## CONCLUSION

Overall, this study provides new findings highlighting the importance of biophysical extracellular signals in cortex development. It describes a novel tool to explore under both physiological and pathological contexts, the two main migratory hehaviors displayed by the inhibitory cortical interneurons. The resulting impact of this research might be multiple, for physiological and pathological conditions, where such a tool can provide new insights into the etiology of neurodevelopmental syndromes.

## Acknowledgements

This work was supported by INSERM, “Institut Pierre-Gilles de Gennes” (“Investissements d’avenir” program ANR-10-IDEX-0001-02 PSL and ANR-10-LABX-31, and ANR-10-EQPX-34), Sorbonne University, by fellowships to CL (PhD funding of the French Ministry of Research attributed by the « Interface pour le Vivant » program of Sorbonne University, and fourth year thesis fellowship from the Fondation Pour la Recherche Médicale, FRM: FDT20170437369), and by grants from the french Agence Nationale pour la Recherche (ANR-14-CE13-0018-01 to CM), from the Fondation Jérôme Lejeune (Grant R14108DD to CM), and through European Research Council Advanced Grant No. 321107 “CellO” (CV). We warmly thank M. Piel (Institut Curie, Paris, France) and N. Spassky (Ecole Normale Supérieure, Paris, France) for advice, our colleagues of the Fer à Moulin Institute and of the Curie Institute for advice and stimulating discussions. We are grateful to J.Masson, M. Darmon, C. Laclef, A. Baffet, V. Grampa, JM Peyrin and JL Viovy for critical reading of the manuscript, and to Melody Atkins for english revision. We gratefully acknowledge the Imaging plateform facility of the Fer à Moulin Institute for the use of their microscopes, the Imaging Plateform of the Jacques Monod Institute for super resolution microscopy (Paris, UMR7592), the Electron Microscopy Facility of the Paris-Seine Biology Institute (IBPS, Paris) and the Technological Platform of the Pierre-Gilles de Gennes Institute for microfluidics.

## Data availability

The raw/processed data required to reproduce these findings cannot be shared at this time due to technical or time limitations.

---

**Supplimentary figure1:**
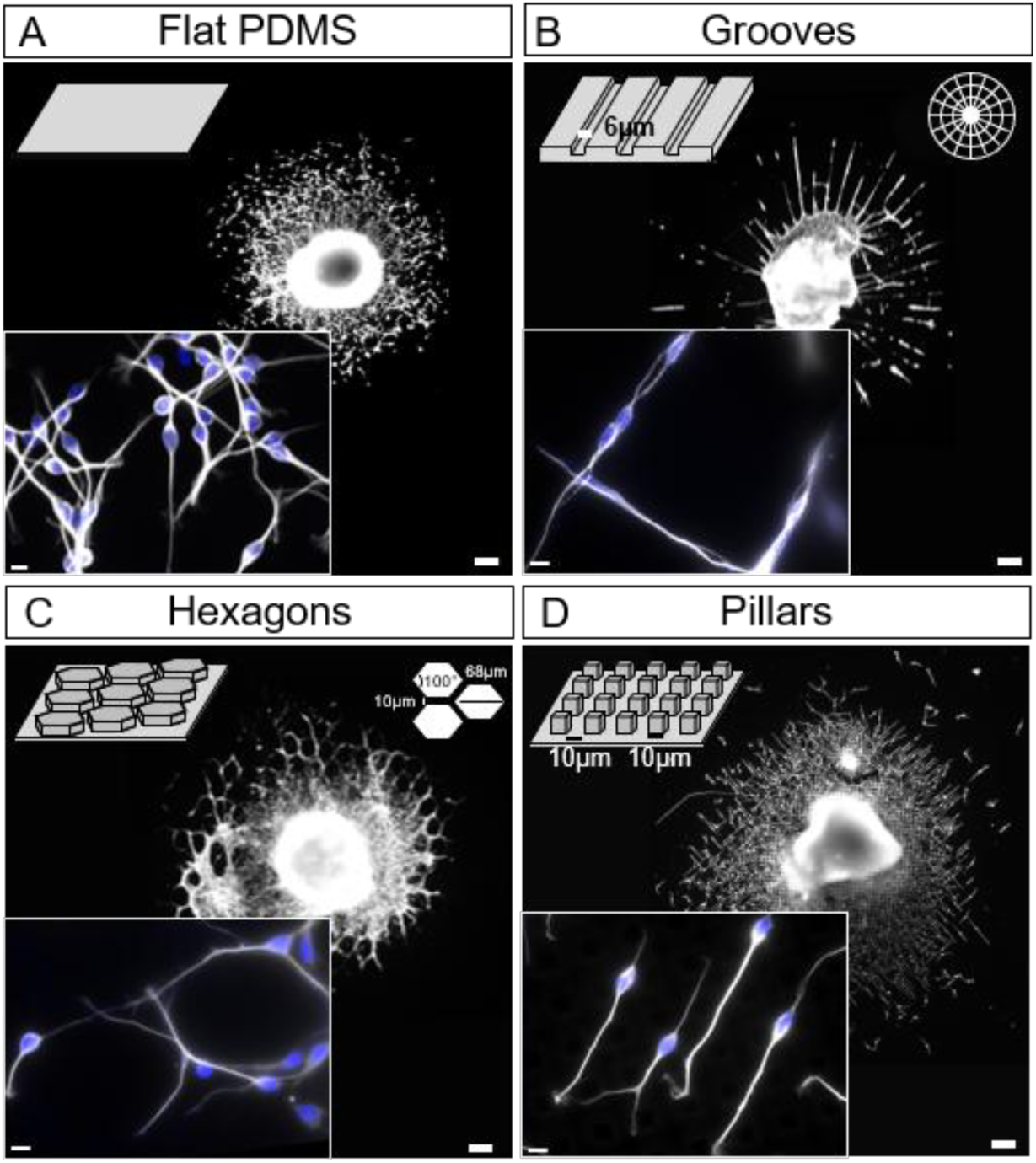
Migration of interneurons on microstructures with different geometries. Explants (white spot in the center of cultures) surrounded by migratory interneurons were fixed after 24 hours in culture. Explants were cultured on either (A) flat PDMS, (B) grooves (width 6 μm, height 10 μm), (C) hexagons (length 68 μm separated by 10μm), or (D) square pillars (10 μm size and height separated by 10 μm). Interneurons are visualized by tyrosinated tubulin immunostaining (white). Inserts show enlarged views of interneurons at the periphery of the migration area (Dapi staining of the nucleus, blue). Scale bars, 100 μm (low magnification picture), 10 μm (inserts).

**Supplimentary figureS2:**
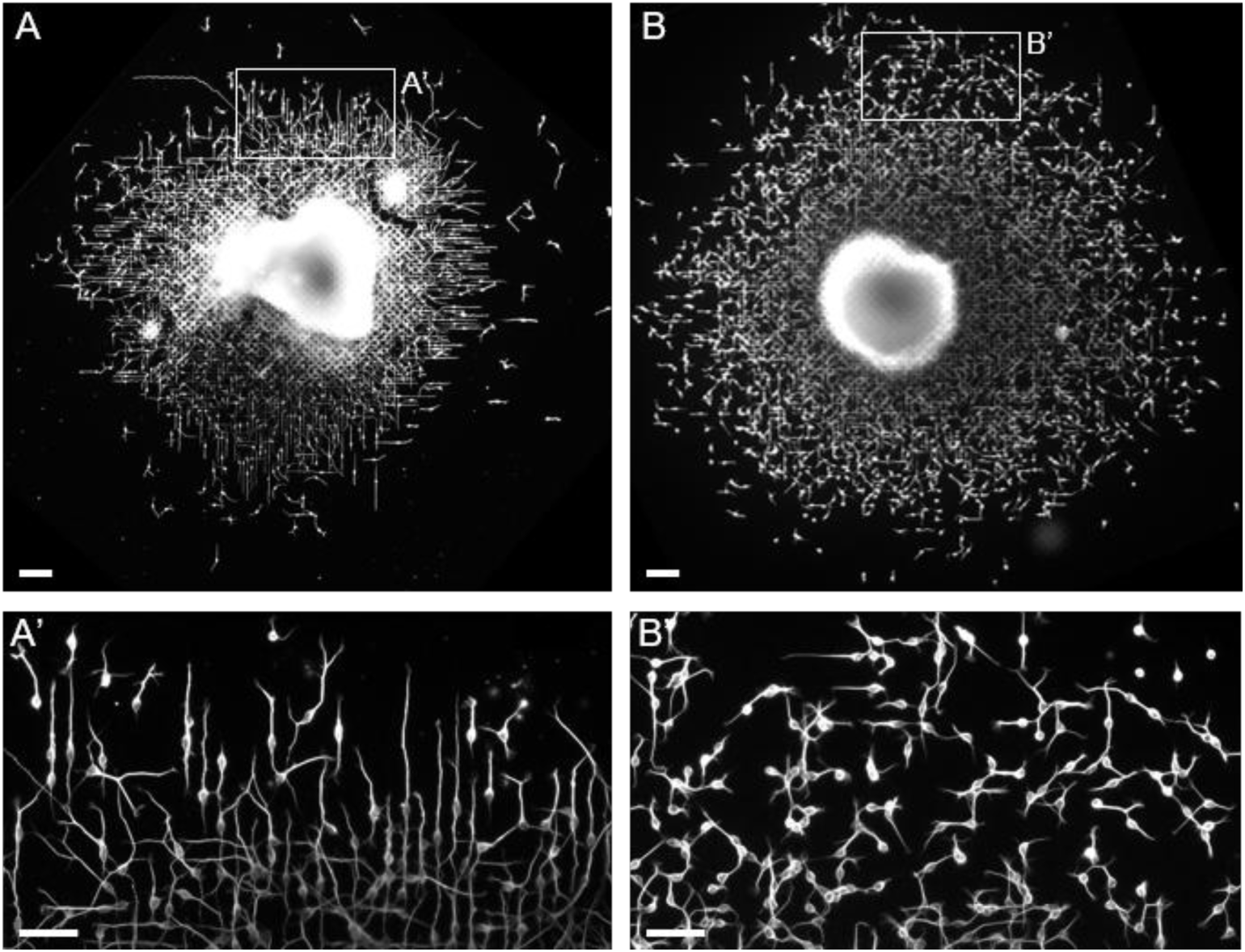
Classification of cultured explants according to the macroscopic organization of migratory interneurons. (A,B) Explants fixed after 24h in culture show two main types of macroscopic organization of interneurons at the periphery of their migration area (A’,B’), either long and aligned (A,A’) or short and non-aligned (B,B’). Scale bars, 100μm (A,B), 50μm (A’, B’)

**Supplimentary figure3:**
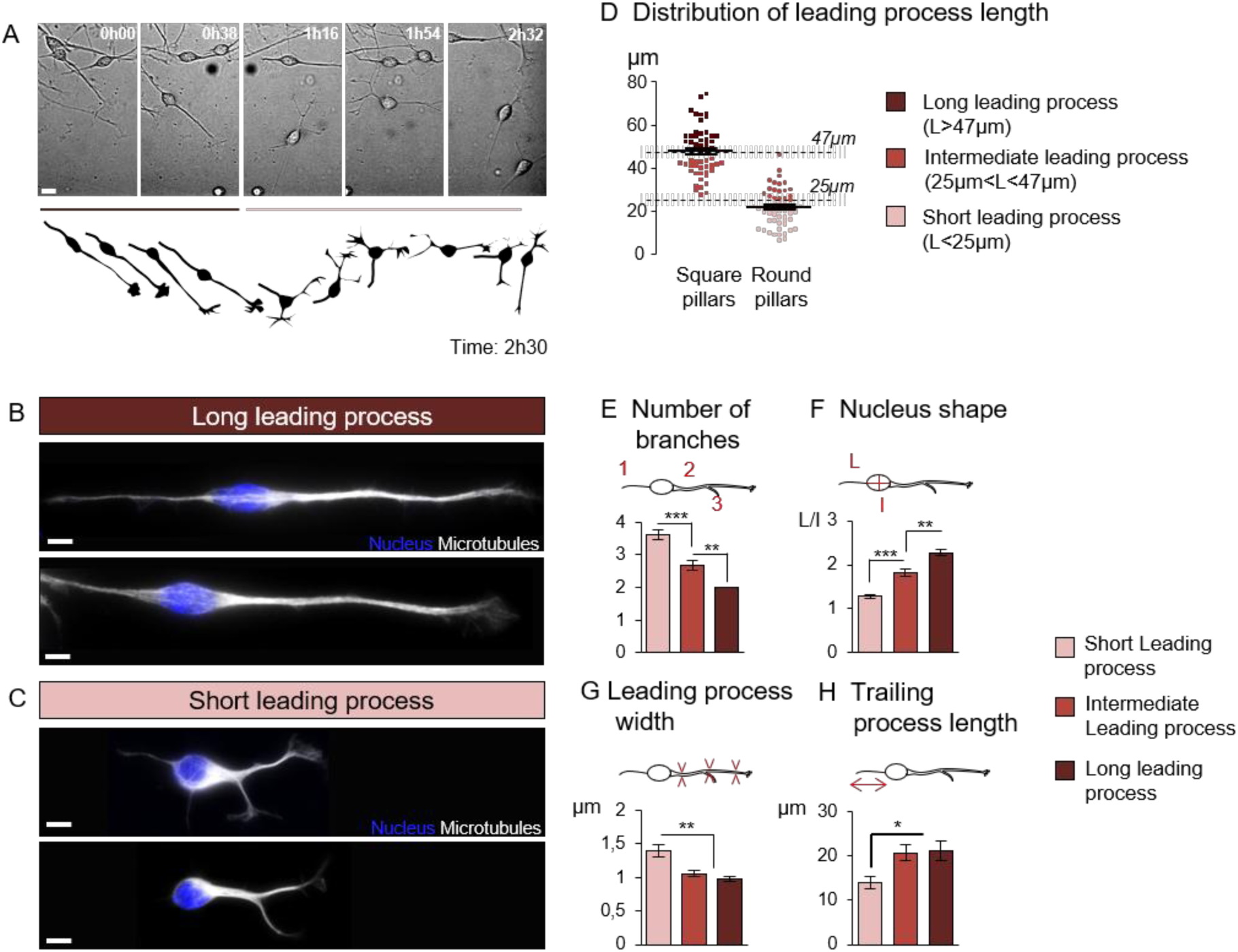
Characterization of two morphological states of migrating interneurons. (A) Frames from the time lapse sequence of an interneuron migrating on a flat surface (top) and drawing of the same sequence (bottom, total duration 2h30). The interneuron exhibited successively two opposite morphological organizations, long unipolar and then, short multipolar. Scale bar, 10μm. (B,C) High magnification views of tyrosinated tubulin (white) and Dapi (blue) labeled interneurons exhibiting those two morphologies. Scale bars, 5μm. (D) Distribution of leading process lengths in interneurons migrating on either square or round pillars (55 long and unbranched cells from 6 cultures on square pillars and 56 short and branched cells from 4 cultures on round pillars shows that interneurons with the longest non-branched leading process were observed on the square pillars, and multipolar interneurons with the shortest processes, on the round pillars. Intermediate lengths were observed on both substrates. (E-H) Interneurons migrating on pillared surfaces were distributed in 3 classes according to the length of their leading process, whatever their morphology (branched, non-branched) and the shape of the pillars (28 cells with a long leading process (6 cultures), 46 cells with intermediate lengths (10 cultures), 37 cells with short leading processes (5 cultures)). In each class, the mean number of branches (E), the nucleus shape (F) defined by the ratio of the long and short axes, the mean width of the leading process (G) (average of 3 measures along the whole length of the LP) and the mean trailing process length (H) were measured.Significance of differences was assessed using a nonparametric ANOVA, Dunn’s post test (***p<0.0001 in E, F, **p=0.0004 in G, *p=0,0069 in H). Error bars represent mean ± SEM.

**Supplimentary figure4:**
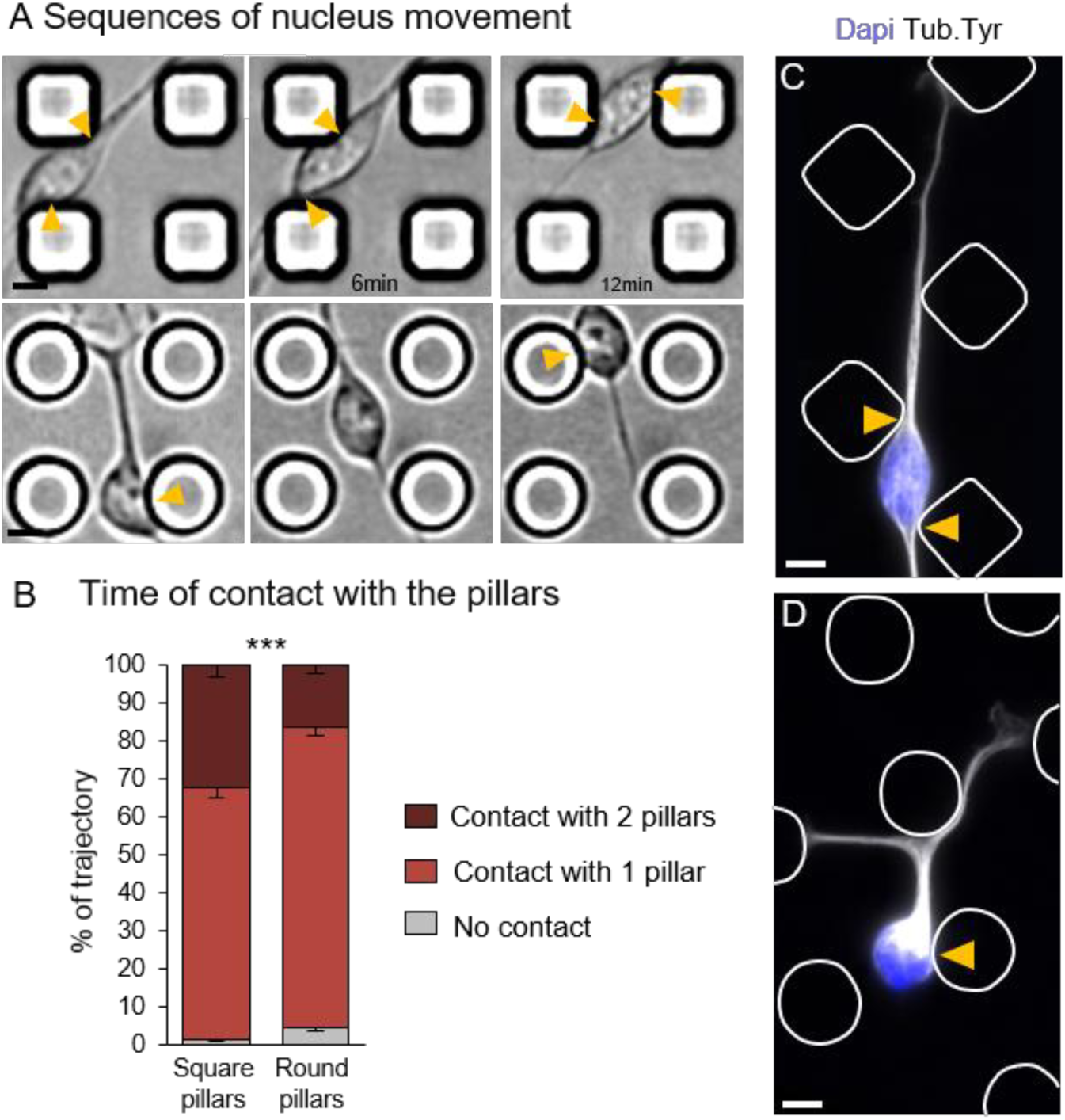
Cell body interaction with the microstructures. (A) Time lapse sequences recorded with a phase contrast microscope showing somal movements in migrating interneurons (top, on square pillars, bottom, on round pillars). Yellow arrows indicate contacts between the cell body and the pillars. Scale bars, 10μm. (B) The time of contact of the cell body with no, one or two pillars is noted on each frame and expressed as a percentage of the total trajectory duration. Contact durations significantly differed on square pillars (30 cells from 6 cultures)and round pillars (30 cells from 5 cultures) (Chi2 test, ***, p<0.0001).Error bars represent mean–SEM. (C,D) Immunostaining of tyrosinated tubulin (white) and Dapi staining (blue) in representative interneurons that migrated on square (C) and round (D) pillars. Pillars were drawn from phase contrast pictures. Contacts between the cell body and the pillars are indicated by arrow heads (yellow). Scale bars, 5μm.

**Supplimentary figure5:**
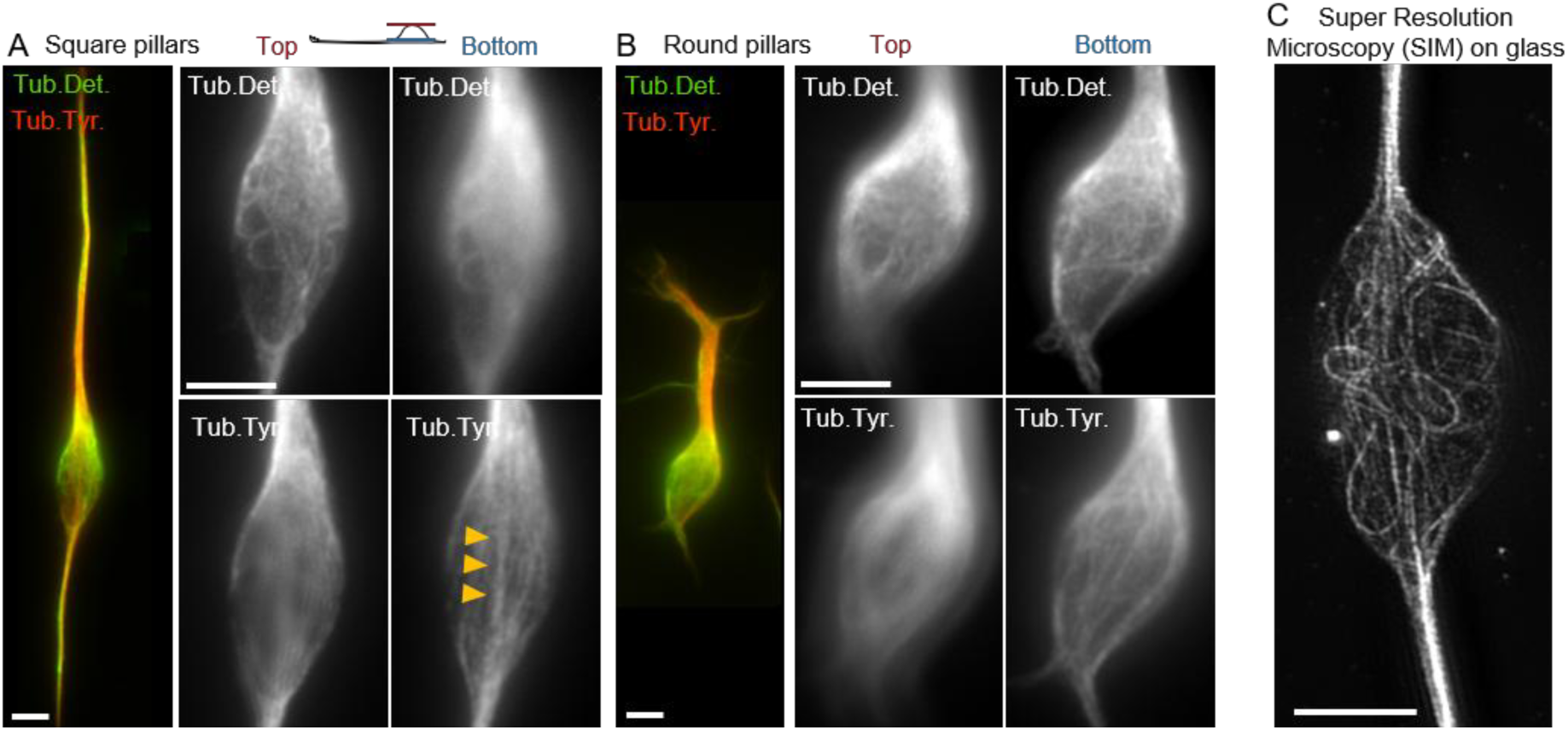
Microtubule network architecture in the cell body. (A,B) Interneurons were immunostained with antibodies against tyrosinated tubulin (dynamic microtubules, red) and detyrosinated tubulin (stable microtubules, green). Panels show maximum projections of epifluorescence microscopy pictures acquired at the top (left) or at the bottom (against the substrate,right) of the cell body. (C) Super resolution microscopy (SIM) shows the organization of stable microtubules (detyrosinated tubulin) in the cell body of an interneuron that migrated on a flat glass coverslip. Scale bars, 5μm.

## Captions of Movies

**Movie 1 related to Figure 4**: phase contrast recording with a X20 objective (NA, 0.7) of symmetric growth cone splitting in between round pillars. Interval between frames: 115 seconds. Yellow arrows indicate the formation of two equivalent sister branches

**Movie 2 related to Figure 4**: phase contrast recording with a X20 objective (NA, 0.7) of asymmetric growth cone splitting in between square pillars. Interval between frames: 105 seconds. Yellow arrow indicates the formation of a thin lateral branch

**Movies 3, 4, 5 related to Figure 5**: phase contrast recording with a X20 objective (NA, 0.7) of interneuron migration in between square pillars (Movie 4), flat surface (Movie 3) or round pillars (Movie 5). Interval between frames: 115 seconds.

**Movie 6 related to Figure 6**: phase contrast recording with a X20 objective (NA, 0.7) of the growth cone movement in between square pillars. Interval between frames: 115 seconds. Red dots show the leading process tip (LPT) tracked for the analysis.

**Movie 7 related to Figure 6**: phase contrast recording with a X20 objective (NA, 0.7) of the growth cone movement in between round pillars. Interval between frames: 115 seconds. Red dots show the leading process tip (LPT) tracked for the analysis

**Movie 8 related to Figure S4A**: phase contrast recording with a X20 objective (NA, 0.7) of cell body navigation in between square pillars. Interval between frames: 90 seconds. Yellow arrows indicate the contacts between the cell body and the pillars

**Movie 9 related to Figure S4A**: phase contrast recording with a X20 objective (NA, 0.7) of cell body navigation in between round pillars. Interval between frames: 105 seconds. Yellow arrows indicate the contacts between the cell body and the pillars

